# Integrative genomic and transcriptomic analyses uncover regulatory landscape of symbiotic nitrogen fixation in soybean natural population

**DOI:** 10.1101/2025.07.18.665310

**Authors:** Yaling Li, Wanjie Feng, Xilong Feng, Xuantong Liu, Shanmeng Hao, Lijie Lian, Luyao Gao, Ying Shao, Hao Chen, Zhao Chen, Jing Yuan, Liya Qin, Xiaoming Li, Xia Li, Xutong Wang

## Abstract

Symbiotic nitrogen fixation (SNF) is a key trait in legume productivity, yet the genetic and regulatory basis underlying its natural variation remains poorly understood. Here, we integrated genome, transcriptome, and chromatin accessibility data from a soybean diversity panel comprising 380 accessions, including 108 wild and 272 cultivated lines. Genome-wide association studies (GWAS) detected multiple loci for SNF traits but with limited resolution due to polygenic architecture and environmental influences. Independent component analysis (ICA) identified 136 co-expression modules; ten ICs were strongly correlated with SNF phenotypes and enriched in circadian clock components (e.g., GmLHY1a/b), lipid metabolism, or defense signaling pathways. Transcriptome-wide association studies (TWAS) linked 1,453, 806, and 178 genes to NFP, NW, and NFE traits, respectively. Among TWAS hits, 185 transcription factors were identified, with 39.0% overlapping selective sweeps, suggesting regulatory evolution under domestication. To further dissect expression regulation, we performed eQTL mapping and detected 4,654 significant eQTLs, including 1,241 local (cis), 2,505 distal (trans), and 908 mixed. By integrating ATAC-seq data from sorted nodule nuclei, we found that eQTLs, particularly local eQTLs, are significantly enriched within open chromatin regions, indicating their regulatory potential. Notably, we validated the circadian clock gene GmLHY1b as a negative regulator of nodulation using CRISPR mutagenesis and CUT&Tag. Our integrative study provides comprehensive genomic and transcriptomic resources from a diverse soybean population, offering novel insights into SNF regulatory networks and a valuable foundation for future SNF research and soybean improvement.

## Introduction

Symbiotic nitrogen fixation (SNF), carried out by rhizobia within legume root nodules, provides approximately 40-70% of the nitrogen required to produce the high-protein seeds that have made soybean (*Glycine max*) the leading plant-based source of dietary protein and the second-largest source of vegetable oil globally (Ciampitti et al., 2021). By converting atmospheric N₂ into bioavailable ammonium, SNF significantly reduces dependence on synthetic nitrogen fertilizers, decreasing production costs and mitigating greenhouse gas emissions, while also enriching soil nitrogen for subsequent crops (Schipanski et al., 2010). Field experiments have demonstrated that even moderate genetic enhancements in nodulation and nitrogen fixation efficiency can deliver measurable yield gains. For instance, soybean mutants such as *ric1a/ric2a*, which produce moderately more nodules without excessive carbon allocation, accumulate greater shoot nitrogen and higher seed yields (Zhong et al., 2024). Consequently, improving SNF efficiency represents a compelling strategy for simultaneously boosting soybean productivity and agricultural sustainability.

Over recent decades, extensive molecular investigations have elucidated key genetic components governing various stages of legume-rhizobia symbiosis, from rhizobial recognition and infection to nodule development, nitrogen fixation, and senescence (Roy et al., 2020). To date, more than one hundred symbiotic genes have been characterized, including LysM-domain receptors (NFR1/NFR5) for Nod-factor perception (Hansen et al., 2024), receptor kinases such as SYMRK/NORK for signal transduction (Chen et al., 2012), calcium-dependent kinase CCaMK and its transcriptional activator CYCLOPS for nodulation gene activation (Singh et al., 2014), and regulators like NSP1/NSP2, ERN1, ENOD40, and CLE peptides for infection-thread and nodule morphogenesis(Crespi et al., 1994; Eckardt, 2009; Mortier et al., 2010; Kawaharada et al., 2017). In mature nodules, proteins such as plant and bacterial hemoglobins, iron transporters (MtVTL8/LjSEN1/GmVTL1a)(Liu et al., 2020; Cai et al., 2024), ureide biosynthesis enzymes, and antioxidant systems sustain nitrogenase activity under micro-aerobic conditions. In contrast, research on the genetic regulation of nitrogen fixation efficiency within mature nodules and nodule senescence remains comparatively limited. Furthermore, little is known about the genetic underpinnings of natural variation in SNF efficiency within soybean germplasm, creating a critical knowledge gap that hampers the exploitation of genetic diversity for SNF improvement.

SNF-related phenotypes, including nodule number (NN), nodule weight (NW), nitrogen fixation activity per plant (NFP), and nitrogen-fixation efficiency normalized by nodule biomass (NFE), and measured leaf chlorophyll content (SPAD), represent complex quantitative traits influenced by multiple small-effect loci and environmental conditions such as soil nitrogen levels, temperature, and rhizobial strains. This complexity limits the effectiveness of traditional genome-wide association studies (GWAS), which have proven highly successful for simpler traits governed by major genes or few loci, such as flowering time (*E1*)(Dong et al., 2022), plant architecture (*Dt1*, *Dt2*) (Tian et al., 2010; Liang et al., 2022), and pod shattering (*Pdh1*)(Funatsuki et al., 2014; Liu et al., 2025). GWAS typically identifies only a limited number of major-effect QTLs, leaving many minor-effect variants unresolved. Moreover, the presence of multiple functional alleles at individual loci can further diminish GWAS statistical power. Additionally, GWAS outcomes are inherently correlative, linking phenotypic variation to broad genomic regions without directly pinpointing causal genes or variants, thus necessitating extensive functional validation. To address these limitations, transcriptome-wide association studies (TWAS) have emerged as a powerful alternative (Ming et al., 2023). TWAS integrates expression quantitative trait loci (eQTL) data with phenotypic variation, treating gene expression as an intermediate phenotype to better identify causal genes (Li et al., 2024). TWAS has successfully dissected complex agronomic traits in multiple crop species, including soybean, where integrated GWAS and TWAS have identified and validated key regulators of seed traits. The effectiveness of TWAS depends significantly on transcriptome datasets from tissues directly relevant to the targeted phenotype (Tang et al., 2021). Given that mature nodules are the primary sites of SNF activity, population-scale transcriptomes derived from mature nodules are ideal for conducting TWAS of SNF traits. Coupling TWAS with high-resolution eQTL mapping and co-localization analysis can further enhance causal inference and regulatory insight.

Modern cultivated soybean originated from wild soybean (*Glycine soja*) in East Asia approximately 6,000-9,000 years ago (Sedivy et al., 2017). Genomic studies have revealed numerous domestication-related selective sweeps affecting traits such as pod shattering, seed dormancy (Wang et al., 2018), seed coat shininess (Zhang et al., 2018), and growth habit (Tian et al., 2010). However, evolutionary changes in SNF-related traits during domestication remain poorly understood. While modern agricultural practices involving intensive nitrogen fertilization might have reduced selective pressure for high SNF efficiency, wild soybeans, typically growing in nutrient-poor environments, may harbor genetic variants enhancing nodule efficiency or conversely altering resource allocation. Comparative analyses between wild and cultivated germplasm thus provide valuable opportunities for identifying beneficial genetic variation that has been lost or inadvertently selected during domestication.

To bridge these knowledge gaps, we assembled a diversity panel of 380 soybean accessions, comprising 108 wild and 272 cultivated lines representing all major soybean-growing regions in China. We conducted phenotyping for multiple SNF-related traits under uniform inoculation conditions, performed deep whole-genome resequencing to generate high-density SNP data, and collected transcriptomic data from mature nodules across the entire population. By integrating comprehensive genetic, transcriptomic, and phenotypic datasets, our study provides an unprecedented dissection of the genetic and regulatory landscapes underlying natural variation in soybean SNF. This work not only clarifies how domestication has shaped SNF traits but also identifies novel genes and regulatory networks that can be leveraged for genetic improvement of soybean SNF, contributing to enhanced yield stability, reduced fertilizer dependency, and more sustainable agricultural practices.

## Results and Discussion

### Divergent genetic structure and SNF-related phenotypic variation in soybean

To dissect the genetic basis of natural variation in symbiotic nitrogen fixation (SNF), we assembled a diversity panel of 380 accessions that included 108 wild soybean (WS, *Glycine soja*) and 272 cultivated soybean (CS, *Glycine max*) lines **(Table S1)**. The collection spans all major soybean–growing regions, capturing broad ecological and domestication gradients **(Figure 1A)**. Specifically, 77 accessions originate from the Northern Spring Region (NSR), 124 from the Huang–Huai–Hai Region (HHHR), and 14 from the South–Central Multiple–Cropping Region (SMCR) **(Figure S1A)**. Whole-genome resequencing at an average depth of approximately 15× produced 6.71 million high-quality biallelic SNPs (minor-allele frequency ≥ 0.05) after stringent filtering. Population-structure analysis resolved two principal clusters separating WS from CS accessions **(Figure 1B)**. When values of *K* greater than 2 were tested, the cultivated cluster split into several sub-clusters, whereas the wild accessions consistently formed a single, homogeneous component **(Figure 1B)**. This pattern indicates that domesticated soybean has undergone lineage-specific differentiation, probably driven by region-specific breeding and selection, whereas the gene pool in WS has remained comparatively uniform. Consistent with the clustering revealed by the population–structure analysis, A SNP-based maximum-likelihood phylogeny corroborated the deep split between the two lineages **(Figure 1C)**. Selective-sweep analysis with the cross-population composite likelihood ratio (XP-CLR) method identified 4,782 genomic intervals that show strong signals between WS and CS, marking likely targets of domestication **(Figure 1D)**. To connect these genomic signals with functional differences, we quantified five SNF-related traits including RW, NW, NN, NFP, and NFE, and SPAD at 28 days post-inoculation (DPI), which corresponds to the V4 vegetative stage of soybean development (**Table S2**). RW is significantly positively correlated with NN (*r* = 0.675, P < 0.001). Likewise, RW shows a strong positive correlation with NW (*r* = 0.961, P < 0.001; **Figure S1B**). Across the three agro–ecological regions, none of the six SNF–related traits differed significantly, most likely because all accessions were inoculated with the same rhizobial strain (*Bradyrhizobium diazoefficiens* USDA110) under uniform greenhouse conditions. The absence of regional differences under a single–strain inoculation implies that, in the field, the composition of indigenous rhizobia communities could be a key determinant of nitrogen–fixation capacity across agro–ecological zones. However, all traits differed significantly between wild and cultivated groups (*P* < 0.01; **Figure 1E**). Cultivated accessions had larger root systems and more nodules (higher RW, NW, and NN), resulting in greater NFP. By contrast, wild soybeans exhibited higher NFE, indicating greater metabolic activity per unit nodule mass. Divergence in SPAD values suggests that photosynthetic capacity has evolved in concert with root symbiotic traits during domestication. Together, these results show that selection for agronomic performance has reshaped both the genetic architecture and the physiological balance between nodule quantity and efficiency in soybean domestication.

**Figure 1.**
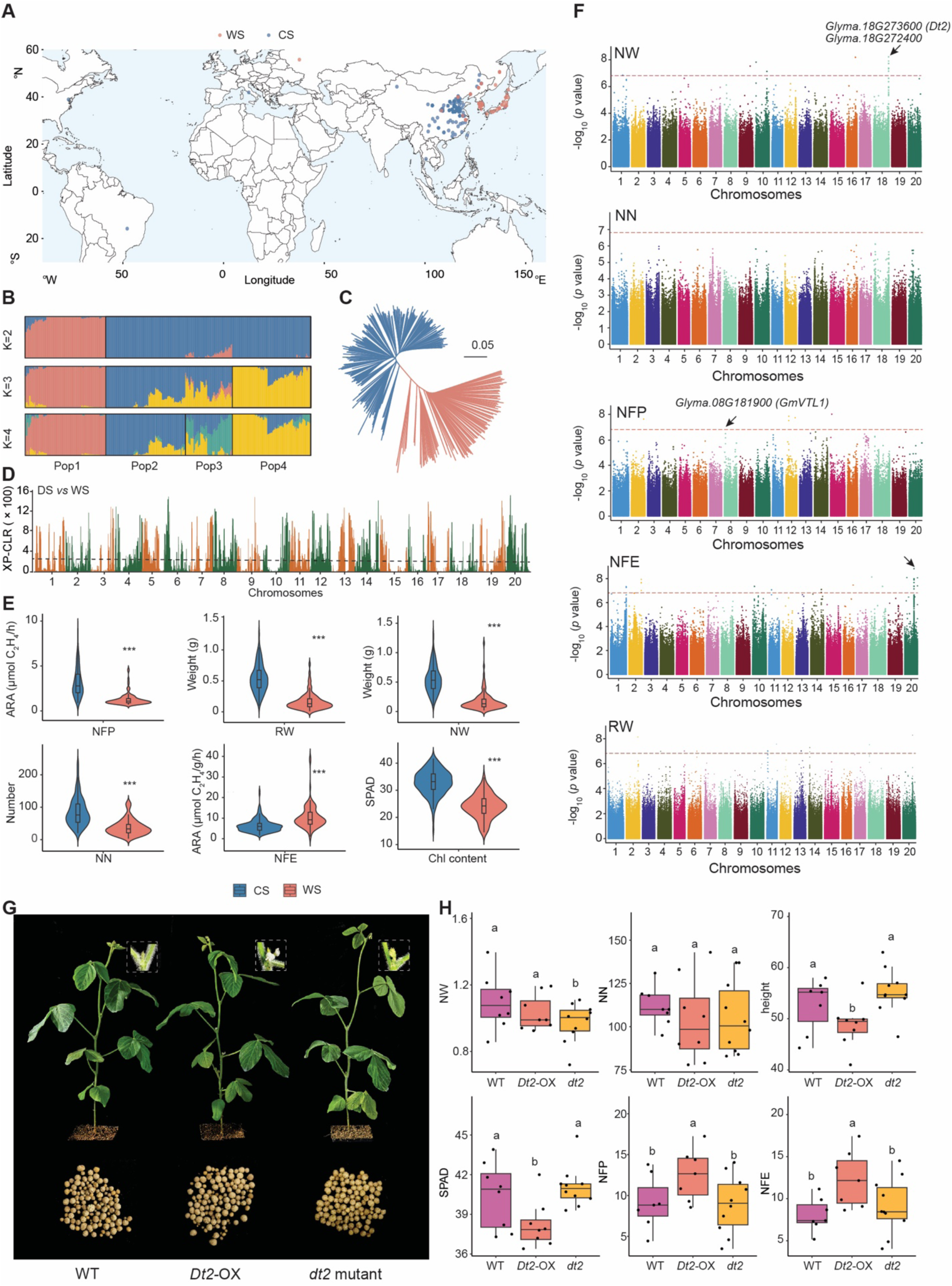
Genetic, phenotypic, and functional analyses of nodulation traits in soybean. **(A)** Geographic distribution of the 380 soybean accessions used in this study, including both cultivated (CS, blue) and wild (WS, red) lines. **(B)** Genomic population structure inferred from genome-wide SNPs using ADMIXTURE (K = 3), with each vertical bar representing one accession. **(C)** Neighbor-joining (NJ) tree constructed from genome-wide genetic distances, illustrating the divergence between cultivated (blue) and wild (red) accessions. **(D)** Genomic regions under selection during domestication, identified using XP-CLR. Regions with XP-CLR scores in the top 10% were considered candidate selective sweeps. **(E)** Phenotypic comparisons between cultivated (blue) and wild (red) accessions for six traits: nitrogen fixation per plant (NFP), root weight (RW), nodule weight (NW), nodule number (NN), nodule nitrogen fixation efficiency (NFE), and chlorophyll content (Chl content, SPAD). Asterisks indicate levels of statistical significance based on Student’s *t*-test (\*\*\**p* < 0.001). **(F)** Manhattan plots of GWAS results for five traits (NW, NN, NFP, NFE, and RW), showing the genomic locations of significantly associated SNPs across the genome. The horizontal dashed line indicates the genome-wide significance threshold (−log10(p) > 6.82). **(G)** Representative shoot and nodule phenotypes of wild type (WT), *Dt2* overexpression (OE), and *Dt2* knockout (KO) plants at 28 days post inoculation. Enlarged views of floral structures are shown to illustrate differences in flowering stage among genotypes. **(H)** Quantification of phenotypic differences among wild type (WT), *Dt2* overexpression (OE), and *Dt2* knockout (KO) plants, including NW, NN, plant height, chlorophyll content (SPAD), NFP, and NFE. Statistical significance was determined using Student’s t-test. Different letters indicate significant differences between groups (p < 0.05).

### GWAS shows limited power to pinpoint key genes controlling SNF traits

To uncover genetic loci associated with phenotypic variation in SNF, we conducted GWAS for six aforementioned traits. We employed the EMMAX linear mixed model to correct for both population structure and relatedness. Significant association signals were identified for four out of the six traits **(Figure 1F)**. For SPAD, a single locus on chromosome 20 surpassed the significance threshold, but no strong candidate gene could be inferred **(Figure S2)**. No loci reached significance for NN or RW in our analysis **(Figure 1F)**. Zhang et al. previously identified *GmNNL1*, an R gene that modulates nodule number, through GWAS in a different soybean panel (Zhang et al., 2021). The absence of NN associations in our study may reflect contrasts in population composition as well as the high environmental sensitivity of NN.

For NW, the strongest association signals lay near *Glyma.18G272400* and *Glyma.18G273600*. *Glyma.18G272400* encodes a B12D–domain protein that is specifically expressed in the uninfected cells of root nodules. *Glyma.18G273600*, better known as *Dt2*, is a key regulator of soybean growth habit. Through its influence on shoot architecture and organ development, *Dt2* may also modulate below–ground traits such as nodule biomass (Liang et al., 2022). To investigate the role of Dt2 in SNF, we compared *Dt2* overexpression (*Dt2*-OX) and *dt2* knockout mutants in the *Williams 82* genetic background. At the R1 developmental stage, *Dt2*-OX plants exhibited earlier flowering and reduced plant height compared to both WT and *dt2* mutants, consistent with the known function of *Dt2* in regulating flowering time and growth habit **(Figure 1G & 1H)** (Ping et al., 2014). However, we did not observe significant differences in NN among *Dt2*-OX, *dt2* mutants, and WT plants **(Figure 1G & 1H)**. Interestingly, the *dt2* mutant showed a reduction in NW compared to WT and *Dt2*-OX, while the latter two lines did not differ significantly in this trait. Although SPAD values were lower in *Dt2*-OX plants, both NFP and NFE were significantly higher than in WT and *dt2* mutant lines **(Figure 1H)**. Overall, the *dt2* mutant did not exhibit pronounced variation in SNF-related traits, possibly due to the low endogenous expression of *Dt2* in the Williams 82 background—also known as the *dt2* genotype. This may limit the phenotypic impact of the *dt2* knockout, thereby attenuating the observable effects of *Dt2* disruption in this genetic context. Among the candidate genes associated with NFP, we identified *Glyma.08G181900* (*GmVTL1*), a homolog of *MtVTL4*, *MtVTL8* and *LjSEN1*(Brear et al., 2020) **(Figure 1F)**. In *Medicago truncatula*, *MtVTL8* is specifically expressed in nodules and required for symbiosome maturation and nitrogen fixation by mediating iron transport into the symbiosome (Walton et al., 2020; Cai et al., 2024). Similarly, *LjSEN1* in *Lotus japonicus* is essential for bacteroid function, and mutants show defective nitrogen fixation due to iron deficiency (Hakoyama et al., 2012). In soybean, GmVTL1a has been shown to localize to the symbiosome membrane and transport ferrous iron into the symbiosome; its mutation leads to reduced nodule iron content and nitrogenase activity (Liu et al., 2020). Furthermore, GmVTL1a can complement the *LjSEN1* mutant, highlighting its functional conservation across legumes (Brear et al., 2020). These findings support a conserved role of *VTL* genes in legume nodule iron homeostasis, suggesting that *GmVTL1* may play a similar role in soybean.

For NFE, we detected a prominent association peak on chromosome 20. Although the lead SNP does not fall within a functionally characterized gene, the surrounding linkage–disequilibrium (LD) block contains six annotated genes (**Figure S3A**). Transcriptome comparisons between CS and WS accessions showed that four of these genes, *Glyma.20G103400*, *Glyma.20G103500*, *Glyma.20G103700*, and *Glyma.20G103800*, are expressed at significantly higher levels in CS lines **(Figure S3B)**. The expression divergence may reflect domestication–driven regulatory changes that contribute to variation in NFE **(Figure 1E)**. Because *Glyma.20G103500* is expressed specifically in nodules, we used a CRISPR/Cas9 hairy-root transformation system to generate knockout lines. Knockout roots were smaller and produced fewer nodules than empty–vector controls **(Figure S3C and S3D)**. Phenotyping revealed no change in root weight, but significant reductions in RL, NN, NW, and NFP. Interestingly, NFE increased in the mutants **(Figure S3D)**. These results indicate that *Glyma.20G103500* promotes root growth and nodule formation while acting as a negative regulator of NFE. Together, these results indicate that natural variation in SNF-related traits is shaped by diverse genetic factors, including several loci overlapping with known nodulation and nutrient transport genes. However, the polygenic nature of these traits and the moderate resolution of GWAS highlight the need for integrative expression-based approaches.

### Population-scale transcriptomics reveals expression networks in mature soybean nodules

To characterize the regulatory programmed that underpin symbiotic nitrogen fixation (SNF), we generated RNA–seq profiles from mature nodules harvested 28 days post–inoculation for 360 accessions. After quality control, 42,398 genes that reached at least 1 FPKM in 5% or more of individuals were retained for analysis **(Dataset 1)**. Principal–component analysis showed extensive overlap between WS and CS accessions, indicating that domestication alone does not impose large constitutive shifts on the mature–nodule transcriptome **(Figure2A)**. Instead, expression values formed continuous gradients across individuals, suggesting that inter–individual variability rather than group–based differences dominates this tissue **(Figure2A)**. Such a continuum supports individual–level mapping approaches, including transcriptome–wide association studies (TWAS) and expression quantitative–trait–locus (eQTL) analyses.

We next decomposed the expression matrix with independent component analysis, which resolved 136 statistically independent components (ICs) containing between 11 and 1,193 genes **(Figure 2B, Table S3)**. Gene Ontology enrichment identified 11 ICs with strong functional coherence **(Figure 2C)**. These modules collectively span key biological processes relevant to SNF. For example, in IC113, which is enriched in GO terms related to secondary cell wall biogenesis and cytoskeleton organization, the presence of *GmSPL9d*, a key transcription factor known to regulate shoot branching and nodule formation (Yun et al., 2022), suggests a potential regulatory link between cell fate determination and structural remodeling processes during development. Modules such as IC34 integrate circadian rhythm, hormone signaling, and stress responses. Key regulators here include *GmLHY1a* and its paralogs, core circadian clock components that influence flowering time, drought tolerance (Wang et al., 2021), and potentially nodule function via temporal coordination of metabolism. In addition, IC71, enriched for cell wall remodeling and enzyme regulation, includes genes like *GmNIC1*, which limits nodule number through CLE peptide signaling (Reid et al., 2013). Their co-occurrence suggests that developmental regulation of organ size and nodulation may rely on shared structural or signaling mechanisms. The lipid metabolism-related IC83 contains *GmWRI1b*, an ethylene-responsive factor critical for seed oil biosynthesis (Zheng et al., 2022), pointing to lipid-based signaling or membrane composition as important for nodule function. Lastly, IC116, enriched for carbohydrate transport and iron homeostasis, harbors *GmbHLH57*, implicated in iron regulation (Li et al., 2018b), and *GmFad3c*, an omega-3 fatty acid desaturase (Singh et al., 2011), potentially linking micronutrient supply and membrane fluidity to efficient nitrogen fixation. Together, these gene-anchored functional modules delineate a transcriptional architecture where developmental, metabolic, and environmental pathways converge to regulate mature nodule physiology, laying a foundation for targeted gene-trait association studies.

**Figure 2.**
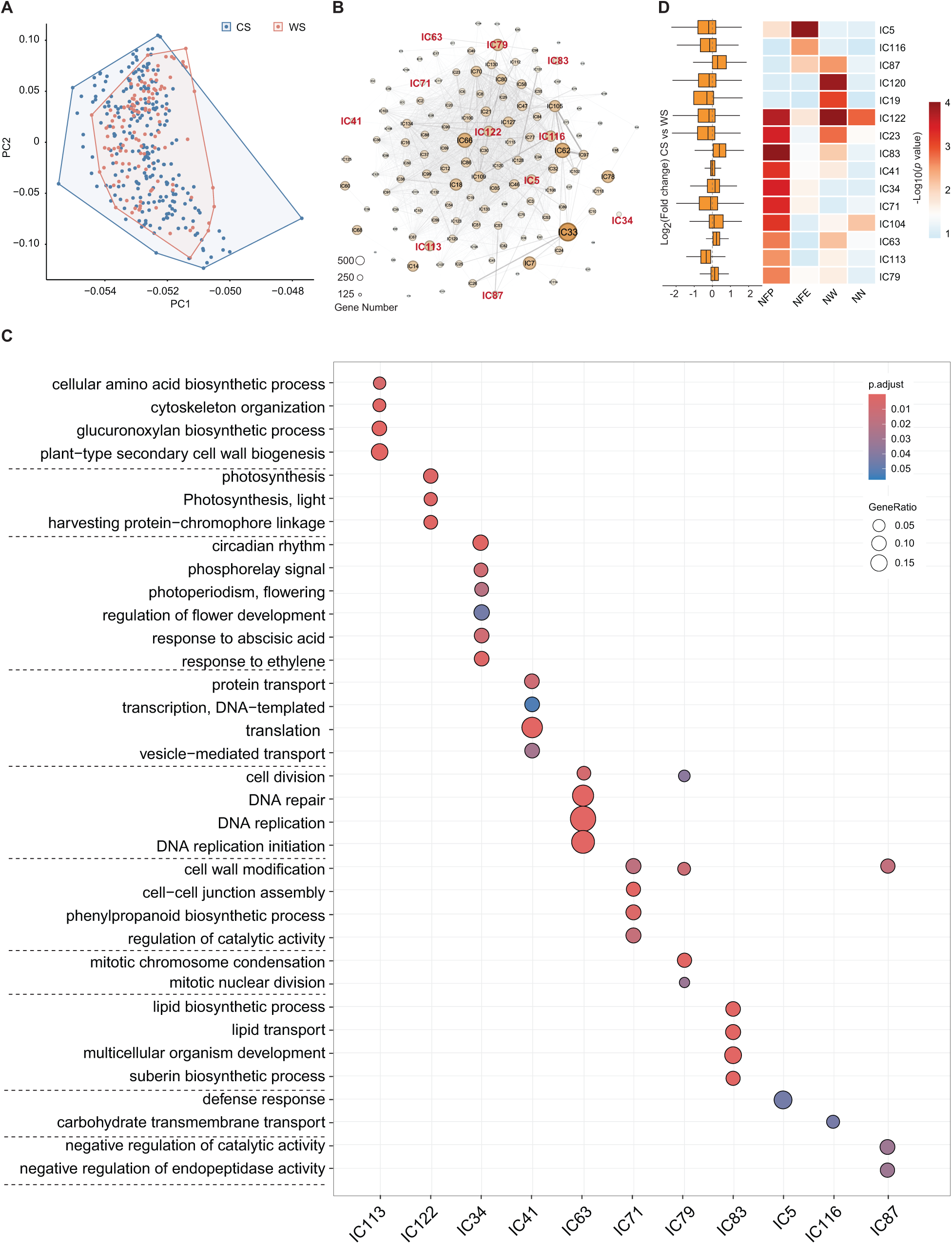
Transcriptomic variation and functional modular structure in soybean. **(A)** Principal component analysis (PCA) of gene expression profiles across 360 soybean accessions. Cultivated accessions are shown in blue, and wild accessions in red. **(B)** Independent component analysis (ICA) of the normalized expression matrix, resolving 136 statistically independent components (ICs). Circle size reflects the number of genes within each component. **(C)** GO enrichment analysis of selected independent components (ICs), including IC5, IC34, IC41, IC63, IC71, IC79, IC83, IC87, IC113, IC116, and IC122. Circle size represents the Gene Ratio (the proportion of genes in each IC associated with a GO term), and color indicates adjusted p-value, ranging from < 0.05 (blue) to < 0.01 (red). **(D)** The associated independent components (ICs) for the four nitrogen fixation traits are shown. The color bar indicates the –log₁₀-transformed p-values. The left panel displays the expression pattern divergence between cultivated soybean (CS) and wild soybean (WS) accessions, measured as the log₂-transformed fold change (CS vs. WS).

To better interpret the biologically significance of gene modules associated with SNF-related phenotypes, we correlated gene expression patterns from ICA-derived modules with trait variation. By examining each trait of NFE, NFP, NW and NN, we aimed to identify functionally coherent transcriptional programs linked to symbiotic processes **(Table S4)**. We found that 15 ICs were associated with at least one trait and exhibited trait-specific enrichment patterns **(Figure 2D)**.

For NFE, genes were strongly enriched in IC5 and IC116, which are associated with defense response and carbohydrate transport, respectively. The overlap between genetic associations and functional enrichment within these modules suggests that pathway involved in Rhizobium interaction and energy allocation may play central roles in regulating NFE. Notably, both modules exhibited higher expression in WS compared to CS **(Figure 2D)**, implying that these genes may contribute to the observed variation in NFE. In contrast to NFE, NFP was associated across ten different modules, with IC83 showing the strongest enrichment **(Figure 2D)**. IC83 is functionally annotated with genes involved in lipid metabolism, pointing to a potential link between membrane remodeling and whole-plant nitrogen fixation outcomes **(Figure 2C)**. These findings highlight the differential regulatory mechanisms underlying local (nodule-level) and systemic (whole-plant) nitrogen fixation traits.

For traits related to nodule development, five IC modules showed significant enrichment for NW, with most exhibiting higher expression levels in WS compared to CS accessions **(Figure 2D)**. The genes within these modules may act as negative regulators of NW, potentially contributing to the reduced NW observed in wild accessions **(Figure 2D)**. Notably, IC122 was one of the most significantly associated modules and was enriched for photosynthesis-related genes **(Figure 2D)**. Although photosynthesis plays a critical role in supplying carbon for symbiotic nitrogen fixation, the negative association between IC122 expression and NW suggests that enhanced carbon assimilation does not necessarily lead to increased nodule biomass **(Figure 2C)**. Rather, it may reflect an imbalance in carbon–nitrogen coupling, where elevated carbon input without synchronized nitrogen assimilation restricts optimal energy allocation to nodule development. This observation underscores the importance of metabolic coordination between source activity (photosynthesis) and sink strength (nodules) in shaping symbiotic organ size. This implies that photosynthesis-derived signals or carbon availability might influence not only nodule growth but also the initiation or proliferation of nodules.

### Nitrogen fixation related genes controlled by the expression regulation in soybean population

To dissect the high associated genes with the phenotype of each trait in NN, NW, NFE, and NFP, we integrated population-level expression data with trait values through TWAS **(Table S5)**. We did not detect any significant gene associate to the NN trait **(Figure 3A)**. This could be due to the use of transcriptomic data from mature nodules at 28 DPI, which may not fully capture the tissue- or stage-specific gene expression patterns underlying NN. A more appropriate sampling strategy targeting relevant developmental stages or tissues might improve the resolution of TWAS for this trait. Except the NN traits, the other three traits we all detected the associated genes, including NFP showing the most associations (1,453 genes), followed by NW (806) and NFE (178) **(Figure 3A, Table S5).** There are 96.6 % (172 out of 178) associate genes present the positive effects between their expression level with the NFE trait while only 3.4% (6 out of 178) present the negative effects **(Figure 3A)**. We also detected some known gene related the nitrogen fixation in the previous studies, such as the NPD and SYP132 genes present the positive effects (Huisman et al., 2016; Pislariu et al., 2019). Besides the known genes, we also found 18 transcription factors (TFs), which may as the key candidate to regulate on NFE trait. 50% (9) TFs were in the selective sweep region, suggest the NFE change in the domesticated process **(Figure 3A, Table S5)**.

**Figure 3.**
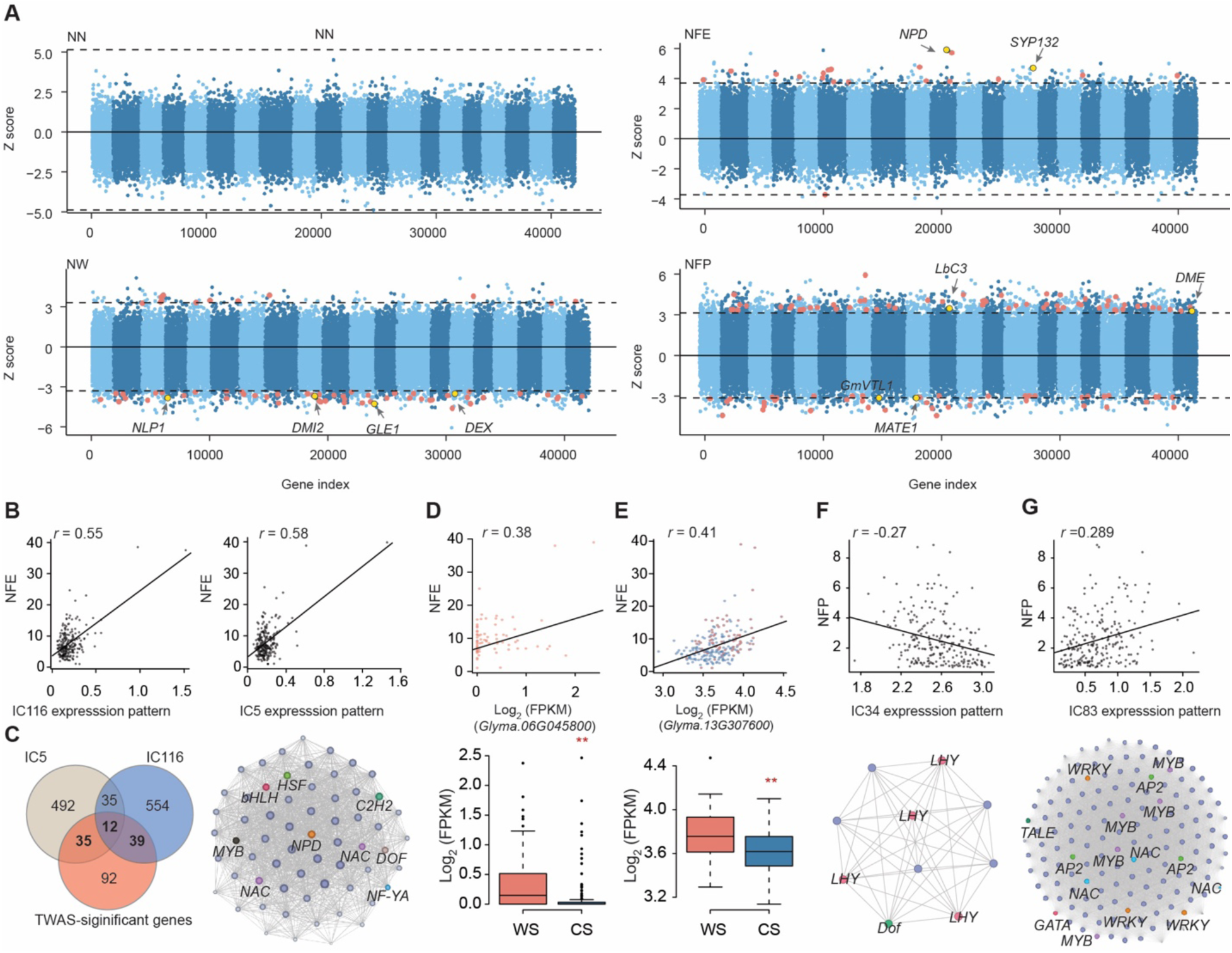
Transcriptome-wide association and expression module analysis reveal regulatory networks underlying nitrogen fixation capacity. **(A)** Transcriptome-wide association study (TWAS) results for four traits: nodule number (NN), nitrogen fixation efficiency (NFE), nodule weight (NW), and nitrogen fixation per plant (NFP). The dotted line indicates the significance threshold. Yellow dots indicate known genes related to symbiotic nitrogen fixation (SNF), while red dots represent transcription factors (TFs). **(B)** Correlation between NFE and the expression patterns of IC116 and IC5. **(C)** Overlap between TWAS-significant genes associated with NFE and genes correlated with IC5 and IC116 (left), and the co-expression network of the overlapping genes (right). **(D)** Correlation between NFE and the expression level of *Glyma.06G045800* (top) and expression levels of this gene in wild (red) and domesticated (blue) soybean accessions (bottom). **(E)** Correlation between NFE and the expression level of *Glyma.13G307600* (top) and expression levels of this gene in wild (red) and domesticated (blue) soybean accessions (bottom). **(F)** Correlation between NFP and the expression pattern of IC34 (top), and the co-expression network of genes within IC34 (bottom). **(G)** Correlation between NFP and the expression pattern of IC83 (top), and the co-expression network of genes within IC83 (bottom).

In total, 83.4% (672 out of 806) genes associated with NW present negative effects of expression on the trait, while only 16.6% (134 out of 806) present positive effects **(Figure 3A)**. This predominance of negatively regulated genes is consistent with the expression patterns observed in the corresponding IC modules **(Figure 2D)**. Among the TWAS-significant genes, four are previously characterized as functionally important in symbiosis or nodule development, including *NLP1*, *DMI2*, *GLE1* and *DEX* **(Figure 3A)**. All four genes exhibited negative effects on NW. *NLP1* is a key transcriptional regulator involved in nitrate signaling and nodule repression under high nitrogen conditions (Luo et al., 2022); *DMI2* encodes a receptor-like kinase essential for early symbiotic signaling (Hogg et al., 2006); *GLE1* is required for mRNA export and essential for normal plant growth and development (Bolger et al., 2008). The negative association of these genes with NW suggests that tight regulation of signaling and metabolic processes may be necessary to control nodule size. In addition, 63 TFs were found to be associated with NW, of which 38.0% (24 TFs) overlapped with selective sweep region, suggesting potential evolutionary selection on transcriptional regulation of nodule weight **(Figure 3A, Table S5)**. In the NFP trait, we detected the large number of associated gene, including 66.8% (970 genes) present the positive effects, including the two known genes, LbC3 and DME (Stougaard et al., 1987; Wang et al., 2024); 33.2% (483 genes) present the negative effects with one known gene, MATE1 (Wang et al., 2016). And 104 TFs were found associated to the NFP trait, 36.5% (38 TFs) include the selective sweep signal **(Figure 3A, Table S5)**. The large associate genes with NFP trait reflected that it is a relative complex trait in soybean, depends on multiple biological processes involved.

We next used NFE as a case study to explore how ICA components may contribute to TWAS-identified associations. Expression levels of IC5 and IC116 were both positively correlated with NFE across accessions **(Figure 3B)**, suggesting these components’ activities reflect or contribute to phenotypic differences. There are 12 genes shared among NFE-TWAS hits and both ICs, with an additional 35 and 39 genes overlapping exclusively with IC5 and IC116, respectively **(Figure 3C)**. Network of these 86 genes recovered nine well-known regulators of nodulation and nitrogen metabolism, such as NAC transcription factors and the Nodule-Specific PLAT Domain Protein (NPD), supporting the biological relevance of these components **(Figure 3C)**. Notably, NAC transcription factors are known to regulate nodule development and senescence (Yu et al., 2023; Wang et al., 2023), potentially by modulating downstream symbiotic genes in response to environmental cues such as nitrate. In addition to the correlation between modules and phenotypes, we identified two individual genes, *Glyma.06G045800* and *Glyma.13G307600*, whose expression was positively correlated with NFE **(Figure 3D)** and significantly upregulated in wild accessions **(Figure 3E)**. These genes may represent previously unrecognized contributors to SNF performance, particularly in wild germplasm.

For the trait NFP, IC83 and IC34 exhibited opposing expression patterns: IC83 was positively correlated with NFP, whereas IC34 showed a negative association **(Figure 3F)**. Functional enrichment analyses revealed distinct biological themes in these two modules **(Figure 2C)**. IC83 was enriched for lipid biosynthetic and transport processes, suberin biosynthesis, and developmental programs related to multicellular organism development, highlighting its potential involvement in maintaining structural and metabolic readiness essential for sustained symbiotic performance **(Figure 2C)**. These processes may be critical for supporting root nodule architecture and function under prolonged nitrogen fixation demands. In contrast, IC34 encompassed genes involved in circadian rhythm, abscisic acid and ethylene signaling, and regulators of photoperiodism and flowering **(Figure 2C)**. The prominence of temporal and hormonal signaling pathways in this module suggests that nodulation in wild germplasm may be tightly coordinated with environmental or developmental timing cues, potentially contributing to the lower NFP observed in WS accessions. Due to the lack of gene overlap between these two ICs, we constructed separate co-expression networks **(Figure 3F & 3G)**. The IC83 network contained seven genes previously implicated in nodulation or nitrogen metabolism, including NAC and AP2 transcription factors. In addition, it featured a broader array of regulatory elements, such as WRKY, MYB, TALE, and GATA transcription factors, which mediate stress responses, metabolic reprogramming, and developmental transitions in legumes (Zhang et al., 2015; Dolgikh et al., 2020; Da Silva et al., 2024) **(Figure 3G)**. The combinatorial presence of these regulators’ underscores IC83 as a potentially central hub supporting effective symbiotic nitrogen fixation. Conversely, the IC34 network featured LHY gene and members of the Dof family, both key regulators of circadian and hormonal responses. Given the known roles of Dof family in plant developmental and stress adaptation (Zou and Sun, 2023), this module likely represents a more responsive or restrictive transcriptional program that modulates nodulation in accordance with environmental conditions **(Figure 3F)**. Together, these results highlight how both modular and gene-level transcriptional architectures shape nitrogen fixation traits in soybean, providing valuable targets for breeding or engineering more sustainable legume-rhizobium symbioses.

### Functional Validation Supports a Negative Role of *GmLHY* Genes in Nodulation

To strengthen the evidence linking the circadian regulators GmLHYs to SNF performance, we quantified all SNF-related traits in CRISPR-generated *Gmlhy* mutants. Relative to the wild type, both *Gmlhy1a* and *Gmlhy1b* mutants showed significant increases in NN and total NW, with *Gmlhy1a* exerting the stronger effect on NN **(Figure 4A & 4B)**. In contrast, overexpression of *GmLHY1b* in transgenic hairy roots resulted in reduced RL, RW, NN and NW **(Figure 4A & 4C)**. These opposing phenotypes support a repressive role for *GmLHY* in the control of nodulation.

**Figure 4.**
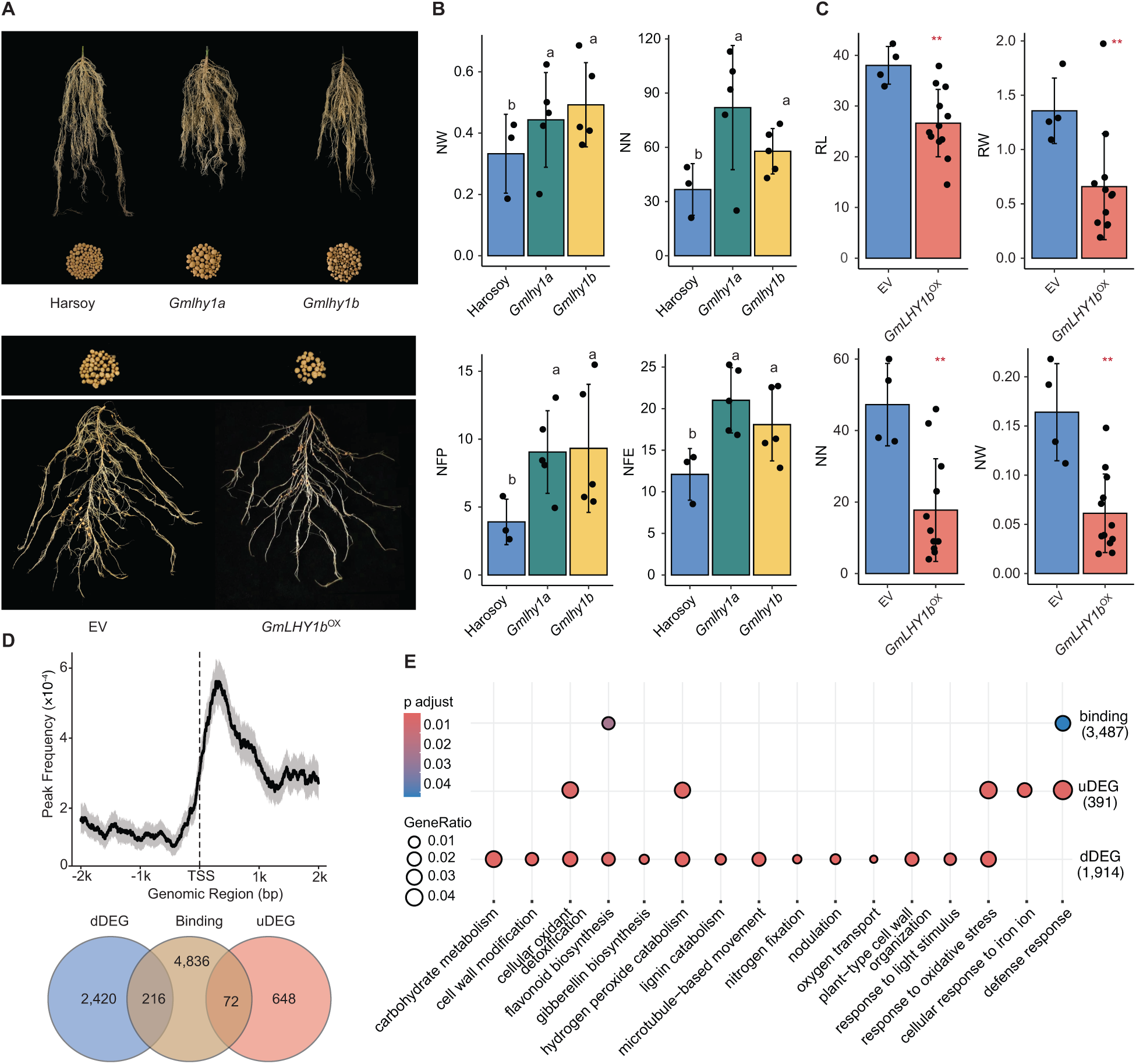
Functional characterization of *GmLHY1a* and *GmLHY1b* in regulating nodulation and root development. **(A)** Representative root and nodule phenotypes of wild-type Harsoy, *GmLHY1a* knockout, and *GmLHY1b* knockout plants (top), and of empty vector control (EV) and *GmLHY1b*-overexpressing (OX) plants (bottom). **(B)** Quantification of phenotypic traits in Harsoy, *GmLHY1a* KO, and *GmLHY1b* KO plants, including nodule weight (NW), nodule number (NN), nitrogen fixation per plant (NFP), and nitrogen fixation efficiency (NFE). Statistical significance was determined using Student’s t-test. Different letters indicate significant differences between groups (p < 0.05). **(C)** Quantification of phenotypic traits in WT (EV) and *GmLHY1b*-overexpressing (OX) plants, including root length (RL), RW, NN, and NW. Asterisks indicate significance levels based on Student’s t-test (*p < 0.01). **(D)** Data quality assessment of CUT&Tag experiments (top) and overlap between differentially expressed genes (DEGs) in OX vs WT and CUT&Tag binding targets (bottom). **(E)** GO enrichment analysis of *GmLHY1b*-OX upregulated genes, downregulated genes, and CUT&Tag binding targets.

By integrating population transcriptomic profiles with chromatin-binding maps from our CUT&Tag experiment, we identified candidate downstream regulatory targets **(Table S6)**. We found that 216 downregulated genes and 72 upregulated genes share promoter regions with GmLHY binding peaks, which cluster near transcription start sites (TSS) **(Figure 4D)**. The concentration of peaks at TSS positions points to direct transcriptional control by GmLHY **(Figure 4D)**. Gene Ontology analysis of the GmLHY1b-responsive gene set showed that the down-regulated targets are strongly enriched for pathways essential to nodulation and symbiosis **(Figure 4E, Table S7)**. Key categories include carbohydrate metabolism, cell-wall modification, detoxification, flavonoid and gibberellin biosynthesis, nitrogen fixation, oxygen transport, and plant-type cell-wall organization et al **(Figure 4E)**. Both up-regulated and down-regulated genes were also enriched for responses to oxidative stress, cellular oxidants, and hydrogen-peroxide catabolism, indicating that GmLHY1b influences redox signaling as well **(Figure 4E)**. GmLHY binding targets, identified from peak-to-gene associations, are strongly enriched for flavonoid-biosynthesis and defense-response terms **(Figure 4E)**. This pattern is consistent with GmLHY repressing early nodulation signals and modulating host–microbe interactions, supporting a model in which the protein functions as a transcriptional repressor that negatively regulates nodulation in soybean.

### eQTL architecture and network analysis uncover local-distal control hubs for soybean nitrogen fixation

To further dissect the regulatory architecture underlying nitrogen fixation efficiency, we conducted genome-wide eQTL mapping to link genetic variants with gene expression variation. This analysis identified 4,654 significant lead associations (threshold: 1.51 × 10⁻⁷), including 1,241 local (*cis*), 2,505 distal (trans), and 908 classified as both **(Figure 5A)**. Clusters of distal eQTLs formed pronounced hotspots, particularly on chromosomes 17 and 18, suggesting the presence of trans-regulatory hubs that may orchestrate broad transcriptional programs across the genome **(Figure 5B, Table S8)**. We identified 1,433 eQTL-eGene pairs located within selective sweep regions (**Table S8**). Interestingly, 685 of these pairs (47.8%) exhibited distal regulatory relationships, suggesting that both regulatory variants and their target genes may have co-evolved during soybean domestication (**Table S8**). Local eQTLs were not only more frequent but also exhibited stronger effect sizes than their distal counterparts **(Figure 5C)**. In total, 2,397 eGenes, approximately 45% of all genes, were associated with at least one significant local eQTL, underscoring the extensive contribution of local regulatory variation **(Figure 5C)**. To complement this analysis, we assessed chromatin accessibility at eQTL loci **(Figure 5D)**. By comparing the observed overlap between eQTLs and open chromatin regions (OCRs) in nodules to a null distribution generated from randomly sampled genomic intervals, we observed a marked enrichment, surpassing the 95th percentile threshold **(Figure 5E, Table S9)**. This signal was particularly strong for local eQTLs, underscoring the functional relevance of accessible chromatin in mediating gene regulatory variation **(Figure 5E)**. We next investigated the genomic context of regulatory loci by examining the positional distribution of eQTLs relative to their eGenes **(Figure 5E)**. Most eQTLs (60.77 %) mapped to intergenic regions, 45.4 % fell within promoter sequences (≤ 2 kb upstream of annotated genes), and only 3.83 % were located inside gene bodies **(Figure 5E)**. Further, we examined whether known genes involved in SNF were present in the eQTL network. Indeed, multiple key regulators were captured. A total of 195 known function genes were identified within the eQTL-derived regulatory networks **(Table S10)**. GmNFR1α (Rj1) and GmNFR5β (Rj6), encoding LysM-type receptor-like kinases that perceive rhizobia Nod factors (Indrasumunar and Gresshoff, 2011; Hayashi et al., 2012), were regulated by both cis- and trans-eQTL. GmNSP1a, a GRAS-domain transcription factor acting downstream in the Nod signaling pathway, was also included. The network also encompassed Rj2/Rfg1, which encodes a TIR-NBS-LRR protein controlling host-Rhizobium compatibility (Yang et al., 2010). Lastly, CHS3C and CHS4A, two key enzymes in flavonoid biosynthesis (Anguraj Vadivel et al., 2018), were represent in the network **(Table S9)**. As flavonoids mediate rhizobia attraction and infection initiation via root exudates, the inclusion of these genes suggests that the regulatory variation in root-microbe signaling is well captured by the eQTL architecture.

**Figure 5.**
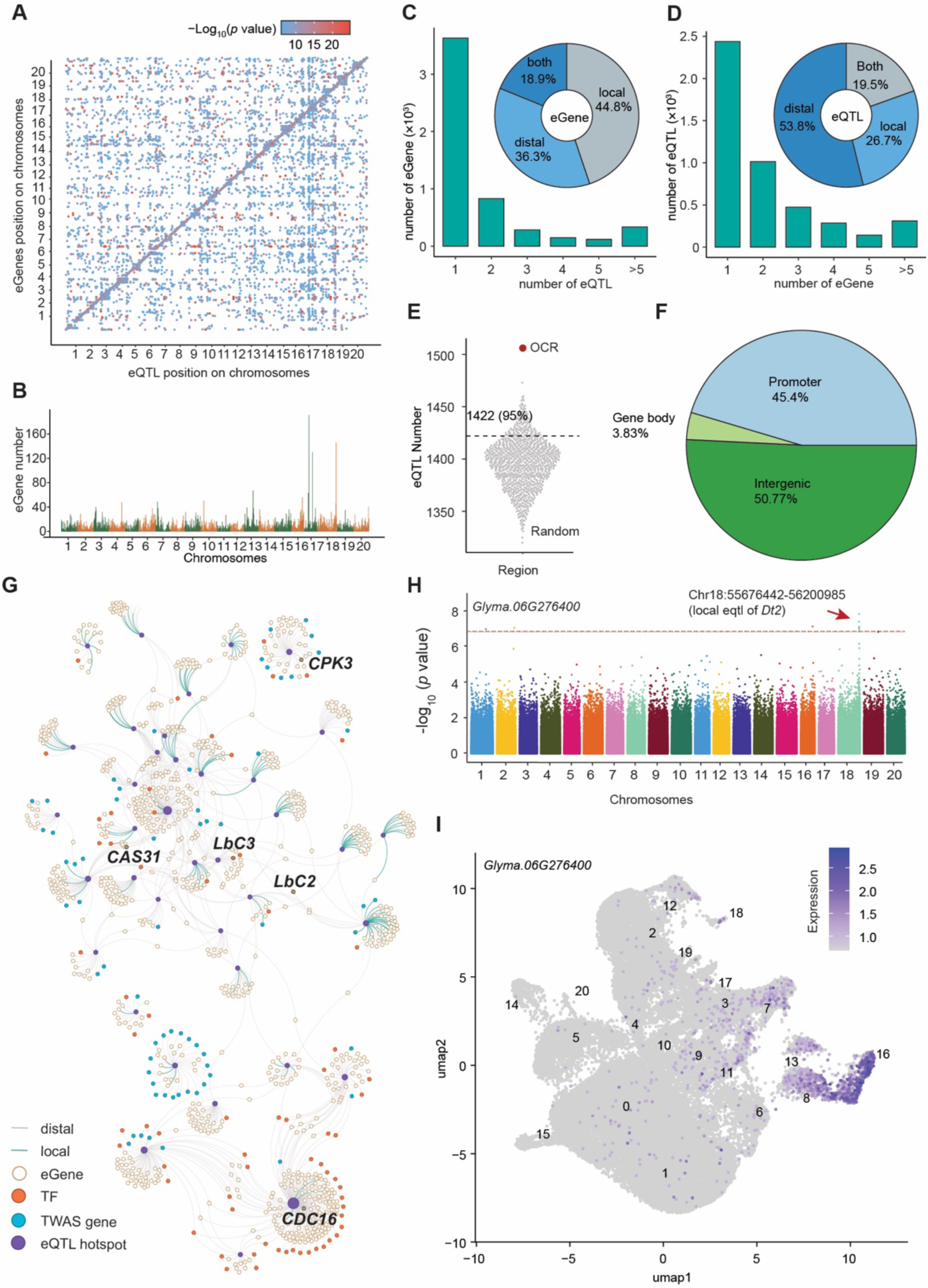
Genome-wide characterization of expression quantitative trait loci (eQTLs) in soybean. **(A)** Genome-wide distribution of eQTL-eGene associations across 20 chromosomes. Each dot represents one eQTL-eGene pair, and color indicates significance (−log₁₀(P)). **(B)** Number of eGenes regulated by eQTLs within non-overlapping 1-Mb windows across the genome. **(C)** Distribution of eGenes by the number of regulating eQTLs, and proportions of eGenes regulated by local versus distal eQTLs. **(D)** distribution of eQTLs by the number of regulated eGenes, and proportions of local versus distal eQTLs. **(E)** Enrichment of eQTLs in open chromatin regions (OCRs), compared with randomly permuted genomic intervals (n = 1,000 permutations). The dashed line represents the 95% confidence interval. **(F)** Genomic context of eQTLs, classified by their relative location to gene features (e.g., promoter, exon, intron, intergenic). **(G)** Genetic regulatory network connecting eQTL hotspots to their downstream eGenes. **(H)** Manhattan plots of eqtl results for the *Glyma.06G276400* gene expression, showing the genomic locations of significantly associated SNPs across the genome. The horizontal dashed line indicates the genome-wide significance threshold (−log10(p) > 6.82). **(I)** Cell-type-specific expression of *Glyma.06G276400* revealed by single-nucleus RNA-seq. The cluster numbers represent distinct cell types, and their correspondence to specific cell identities is illustrated in Figure S4.

Building upon these findings, we constructed a gene regulatory network centered on eQTL hotspots **(Table S8)**. A total of 38 hotspots were identified, many of which exert broad regulatory effects on key genes, including transcription factors and TWAS-significant genes **(Figure 5G**). Notably, most of the hotspots act locally through *cis* regulation, while others regulate a wide range of distant eGenes via trans-acting mechanisms. This contrast highlights the diverse regulatory strategies by which eQTL hotspots shape the transcriptional landscape of soybean nodules. For instance, CPK3, a calcium-dependent protein kinase involved in signal transduction, was located in a distal-only hotspot network that also contained eight TWAS-significant genes and three transcription factors. *LbC2* and *LbC3*, which encode leghemoglobin essential for oxygen transport in nodules (Stougaard et al., 1987; Ahmad et al., 2023), were regulated by two distal eQTL hotspots, one of which simultaneously targeted both genes, suggesting shared upstream control. Similarly, *CAS31*, a stress-responsive gene (Li et al., 2018a), was under distal regulation. *CDC16*, associated with cell division (De Oliveira et al., 2022), was embedded in the largest hotspot network, which was primarily trans-acting but also included three local targets, suggesting that active cell proliferation during nodule development may be partially governed by this regulatory hub. In addition, the network captured several well-characterized genes involved in SNF.

To investigate the regulatory basis of TWAS-identified genes, we overlapped them with eGenes from our eQTL analysis. We found that 355 TWAS genes were also eGenes, with 71.7% regulated through distal eQTLs **(Table S8)**. Further colocalization with GWAS signals revealed a single overlapping eQTL region (Chr18:55676442–56200985) that distally regulates the TWAS gene *Glyma.06G276400*, which is associated with NW **(Figure 5H & 2F)**. Notably, this eQTL also acts locally on *Dt2*, a known developmental regulator. These findings support our earlier hypothesis that Dt2 may also influence symbiotic nitrogen fixation traits. *Glyma.06G276400*, also displayed strong cell-type specificity in nodule single-cell RNA-seq data, being preferentially expressed in rhizobia infection zones **(Figure 5I)**. Such spatially restricted expression points to a potential role in mediating early host-microbe interactions. Together, these results demonstrate that eQTL-informed regulatory networks not only include core SNF genes but also reveals new regulatory candidates, offering a systems-level view of the genetic architecture underlying symbiotic nitrogen fixation in soybean.

Overall, our study provides a comprehensive, systems-level understanding of how natural variation and domestication have shaped the balance between nodule quantity and efficiency in soybean. By connecting genomic variants, gene expression programs, and phenotypic outcomes, we have illuminated the complex genetic architecture underlying biological nitrogen fixation in soybean. The extensive genomic and nodule transcriptomic datasets for 380 accessions generated here are valuable resources for the community. These data can be mined to further dissect legume– rhizobium interactions and will inform breeding or biotechnological strategies to develop soybean varieties with improved nitrogen-fixing performance, reducing reliance on synthetic fertilizers and promoting sustainable agriculture.

Despite the breadth of our integrative analyses, several limitations warrant consideration. First, although our population-scale transcriptomic dataset captures expression variation from mature nodules at 28 days post-inoculation (DPI), it does not encompass the full temporal and spatial dynamics of nodulation. Key regulatory events controlling early infection, nodule initiation, or senescence may have been missed due to stage-specific gene expression that is not represented in our sampling window. Similarly, transcriptomic data were obtained from bulk nodule tissue rather than single-cell or cell-type-resolved profiles, potentially masking regulatory variation in specific nodule zones such as the infection zone or the vascular bundle. Second, while our GWAS, TWAS, and eQTL analyses provide valuable insights into the genetic architecture of SNF-related traits, the resolution of association signals is still constrained by linkage disequilibrium and the polygenic nature of these complex traits. Many candidate genes remain correlative, and functional validation was only performed for a limited subset, such as GmLHY1b. Additional causal relationships remain to be established through targeted genetic or biochemical validation. Third, although we utilized a controlled greenhouse environment with uniform rhizobial inoculation, this does not fully replicate the environmental heterogeneity and microbial diversity encountered in field conditions. Environmental variables such as soil nitrogen content, native rhizobial strains, and abiotic stressors likely modulate SNF outcomes and may interact with the identified regulatory loci. Future studies incorporating field trials and microbiome profiling would enable a more ecologically relevant assessment of SNF trait regulation. Lastly, the current study focuses exclusively on a Chinese soybean diversity panel. Expanding similar integrative analyses to global germplasm collections will help uncover novel alleles and regulatory variants that are absent in Chinese lines but potentially valuable for breeding programs elsewhere. Future efforts should prioritize multi-timepoint, multi-tissue, and single-cell transcriptomic profiling to refine the spatial–temporal resolution of SNF regulation. Moreover, integrating additional layers such as epigenomic, proteomic, and metabolomic data would further elucidate the multilayered regulatory networks governing nitrogen fixation. Ultimately, the integration of high-resolution functional genomics with genome editing and synthetic biology approaches holds promise for translating regulatory insights into improved SNF efficiency in soybean and other legumes.

## Materials and methods

### Plant materials and phenotype collection Measurement of nitrogen fixation traits

For phenotyping, 380 soybean (*Glycine max* and *Glycine soja*) accessions were grown in a greenhouse under controlled conditions of 25 °C, 65% relative humidity, and a 16-hour light/8-hour dark photoperiod. On the fourth day after germination, seedlings were inoculated with *Bradyrhizobium diazoefficiens* strain USDA110. During plant growth, irrigation was carried out alternately with water and nitrogen-deficient BD nutrient solution. Phenotypic evaluation of six traits was conducted at 28 days post inoculation (28 DPI). Measurement of nitrogenase activity was followed the previously described acetylene reduction assay (ARA) system (Yu et al., 2023).

### Whole-Genome Resequencing and Genetic Variation Calling

Genomic DNA was extraction from V1 stage leaves using the CTAB method (Allen et al., 2006), then constructed resequencing libraries using MGIEasy DNA Library Prep Kits. The libraries were sequenced on MGI DNBSEQ-T7 platform for 150 bp paired-end reads. Raw sequencing reads were first assessed for quality using FastQC (0.11.9) (https://www.bioinformatics.babraham.ac.uk/projects/fastqc/), and low-quality bases and adapter sequences were removed using Trim Galore (version 0.6.10) (https://github.com/FelixKrueger/TrimGalore). Clean reads were aligned to the Glycine max reference genome (Williams 82, version a4.v1) using BWA-MEM (version 0.7.17-r1188) (Li, 2013). SAMtools (version 1.18) (Danecek et al., 2021) was used to sort and index the alignments, retaining only properly paired reads with MAPQ ≥ 30. PCR duplicates were marked using Picard MarkDuplicates (version 2.27.4) (Broad Institute, 2019). Variant calling was performed using GATK4 (v4.2.6.1) (McKenna et al., 2010). HaplotypeCaller was run in GVCF mode per sample, followed by joint genotyping with GenotypeGVCFs. SNPs were filtered according to GATK best practices, removing variants with QD < 2.0, QUAL < 30.0, MQ < 40.0, SOR > 3.0, FS > 60.0, ReadPosRankSum < –8.0, and MQRankSum < –12.5. Additional filtering using VCFtools (version 0.1.16) excluded SNPs with minor allele frequency (MAF) < 0.05 and missing genotype rate > 10% (Danecek et al., 2011). After filtering, 6,710,329 high-quality SNPs were retained.

### Population Structure and Selective Sweep Analysis

Population structure was analyzed using genome-wide SNPs with ADMIXTURE (version 1.3.0) (Alexander et al., 2009), with the number of ancestral populations (K) set to 3. The optimal K value was supported by cross-validation error. A neighbor-joining phylogenetic tree was constructed in MEGA 7 based on the same SNP dataset (Kumar et al., 2016), using the p-distance model and 1,000 bootstrap replicates for branch support. To identify genomic regions under selection, XP-CLR (https://github.com/hardingnj/xpclr) analysis was conducted between cultivated and wild soybean populations. A sliding window of 100 kb and a grid size of 10 kb were used. Regions within the top 10% of XP-CLR scores were designated as putative selective sweeps.

### Genome-Wide Association Study (GWAS)

GWAS was conducted for six phenotypic traits (NN, NW, RW, NFE, NFP and SPAD) using the command-line version of EMMAX (Kang et al., 2010), separately for cultivated and wild soybean accessions. A kinship matrix was included to account for cryptic relatedness, and the top three principal components (PCs), calculated using GCTA v1.94.1 (Yang et al., 2011), were included as covariates to control for population structure. Association results from the two groups were combined using METAL with a sample-size-based meta-analysis strategy(Willer et al., 2010). Statistical significance was determined based on the effective number of independent markers (Me = 6,613,265), estimated using GEC v1.0 (Li et al., 2012). Associations with P-values below 1/Me were considered significant.

### RNAseq and Data processing

Total RNA was extracted from nodules at 28 DPI using the FastPure Universal Plant Total RNA Isolation Kit (Vazyme, China) following the manufacturer’s instructions. RNA quality and integrity were assessed using an Agilent 2100 Bioanalyzer (Agilent Technologies, USA). Strand-specific RNA-seq libraries were prepared using the MGIEasy RNA Library Prep Kit (BGI, China) according to the manufacturer’s protocol, then sequenced on the MGI DNBSEQ-T7 platform (BGI, China) to generate 150 bp paired-end reads. Each sample yielded approximately 6 Gb of raw RNA-seq data. After trimming with Trimmomatic (version 0.39) (Bolger et al., 2014), clean reads were aligned to the Glycine max reference genome using HISAT2 (version 2.2.1) (Kim et al., 2019). Gene-level read counts were generated using HTSeq-count (version 2.0.3) (Anders et al., 2015), and expression levels were normalized as FPKM (fragments per kilobase per million mapped reads). Genes with FPKM > 1 in at least 5% of samples were retained. The expression matrix was transformed using log₂ (FPKM + 1) and used for subsequent TWAS and eQTL analysis. To extract latent transcriptional programs, Independent Component Analysis (ICA) was performed using the FastICA algorithm implemented in the scikit-learn (https://scikit-learn.org/stable/index.html) Python package. The number of components was set to 136 based on preliminary dimensionality reduction and model stability. Each independent component (IC) represents a statistically independent gene expression module potentially linked to specific regulatory processes. To assess the enrichment of GO terms and KEGG pathways, we carried out a hypergeometric test using the R package clusterProfiler (version 4.10.1) (Wu et al., 2021).

### H2B-GFP transgenic hairy roots, ATAC-seq and data processing

Since mature nodules contain both plant cells and bacteria, we aimed to selectively isolate plant nuclei by combining a nucleus-specific GFP signal (via an H2B-GFP fusion protein) with DAPI staining. These two fluorescent signals allowed us to efficiently sort plant nuclei using fluorescence-activated cell sorting (FACS). For the generation of H2B-GFP transgenic hairy roots, the full-length coding sequence of histone 2B (H2B; *Glyma.11G141600*) was amplified from *Glycine max* cv. Williams 82 genomic DNA using the following primers:

H2BGm11G141600F: AGAACACGGGGGACTCTAGAATGGCACCAAAGGCAG

H2BGm11G141600R: CCCTTGCTCACCATGGATCCAGAGCTAGTGAACTTGGTGACA

The PCR product was inserted upstream of the GFP coding sequence in the pB35S: GFP-BS2 vector via XbaI and BamHI restriction sites (R0145M and R3136M; New England Biolabs). Transgenic hairy roots expressing Pro35S:H2B-GFP were generated in soybean via *Agrobacterium rhizogenes* strain K599 mediated transformation. Seven days after root emergence, the transgenic roots were inoculated with *B. diazoefficiens* strain USDA110 to induce nodule formation.

At 28 DPI, nodules exhibiting green fluorescence were identified using a handheld fluorescence detector. Approximately 0.4 g of fluorescent nodules were harvested and immediately chopped in 500-1000 μL of ice-cold lysis buffer (20 mM MES, pH 8.0; 10 mM MgCl₂; 20 mM KCl; 3 mM DTT; 400 mM D-sorbitol; 0.4% Triton X-100; 1% BSA). The homogenate was filtered twice through Miracloth, stained with DAPI, and subjected to fluorescence-activated nuclei sorting using a BD FACSAria™ SORP flow cytometer. Excitation wavelengths for DAPI and FITC were set to 358 nm and 488 nm, respectively. A total of 50,000 nuclei were sorted based on size, DAPI signal intensity, and FITC signal, and collected into 400 μL of lysis buffer. Nuclei were pelleted by centrifugation at 1000 × g for 10 min at 4 °C. Tn5 transposition and library construction were performed using the TruePrep DNA Library Prep Kit V2 for Illumina (Vazyme) following the manufacturer’s protocol. Amplified libraries were purified using Vazyme magnetic beads and quantified with a Qubit fluorometer. Libraries were then multiplexed and sequenced on an Illumina NovaSeq 6000 platform to generate paired-end reads.

To ensure high-quality sequencing data, raw reads were trimmed and filtered using Trimmomatic (v0.39) to remove low-quality bases and contaminants (Bolger et al., 2014). Cleaned reads were then aligned to the soybean reference genome (Gmax v4, Phytozome) using bowtie2 (v2.3.5.1) (Langmead and Salzberg, 2012). Reads mapping to multiple genomic locations were filtered out using Samtools (v1.7) (Danecek et al., 2021), and PCR duplicates were removed with Picard tools (v1.112) (Broad Institute, 2019). Open chromatin regions were identified genome-wide using MACS2 (v2.2.7.1) with parameters --nomodel --shift -100 --extsize 200 -B -q 0.05 (Zhang et al., 2008).

### GmLHY1b vector construction, hairy root transformation, CUT & Tag, and data processing

To generate constructs with *GmLHY1b* overexpression, the CDS of *GmLHY1b* (*Glyma.07G048500*) was amplified from the *Glycine max* cv. Williams 82 genomic using the following primers:

07G048500-UTR-F: TGCAGTAACATCATCACCATACC

07G048500-UTR-R: CAATATAGTTACACCGTTGTGGG

LHY1b-cds-F: tagtggatcccccctagaaggcctATGGACGCATACTCCTCCG

LHY1b-cds-R: ttaattaacccgctggtaccAGTCGAAGTCTCCCCTTCCA

Then, the PCR prodcut was cloned into the UBIpro:4×MYC vector between StuI and KpnI restriction sites using the In-Fusion Cloning, and Sanger sequecing was performed to confirm the sequence. For generation of LHY1b-OX transgenic hairy roots, the recombinant vector was delivered into *Agrobacterium rhizogenes* strain K599 by the heat shock method. The *B. diazoefficiens* strain USDA110 inoculation treatment was carried out as previously described.

Nuclei were isolated following the ATAC-seq protocol and CUT & Tag assays were performed using the CUT & Tag Assay Kit (TD904), according to the manufacturer’s instructions. Sequencing reads were aligned to the soybean reference genome (Gmax v4, Phytozome) using bowtie2 (v2.3.4.3) (Langmead and Salzberg, 2012). GmLHY1b binding peaks were identified by MACS2 (fold enrichment > 2, q-value < 0.05) (Zhang et al., 2008). Target genes were assigned using BEDtools (version 2.30.0) (Quinlan and Hall, 2010). The resulting CUT & Tag BAM files were converted to bigwig format with deepTools (v3.5.0) for visualization (Ramírez et al., 2014).

### Transcriptome-Wide Association Study (TWAS) and Expression Quantitative Trait Loci (eQTL) Mapping

TWAS was conducted to link gene expression variation to phenotypic traits. For each of the four traits (NN, NW, NFE, and NFP), TWAS was performed separately in cultivated and wild soybean accessions using the cGWAS.emmax function from the cpgen R package (Kang et al., 2010). The results were combined using METAL (Willer et al., 2010), and significant trait–gene associations were identified based on a false discovery rate (FDR) adjusted *p* value threshold of 0.05.

eQTL mapping was conducted to identify genomic loci regulating gene expression. SNPs were pruned for linkage disequilibrium using PLINK 1.9 (Purcell et al., 2007). Association analysis was carried out separately in cultivated and wild accessions using EMMAX (Kang et al., 2010), incorporating a kinship matrix and the top three PCs (from GCTA) to control for population structure. Results from both groups were combined using METAL (Willer et al., 2010). Significance was assessed using the effective number of independent markers (Me, estimated via GEC v1.0) (Li et al., 2012), and associations with P < 1/Me were considered significant. To define independent signals, significant SNPs within a 1 Mb window of a given gene were grouped, and the most significant SNP (lead SNP) was retained. Lead eQTLs were classified as local (*Cis*) if located within 1 Mb upstream of the transcription start site (TSS) or 1 Mb downstream of the transcription end site (TES); otherwise, they were defined as distal (trans).

## DATA AVAILABILITY

The resequencing, transcriptome, and genotype datasets generated and reported in this study have been deposited in the GSA database under accession number PRJCA043389.

## AUTHOR CONTRIBUTIONS

XW, XL conceived and designed the experiments. XF collected all the germplasm and measure the phenotypes used in this study. XW, YL, WF and SH were responsible for performing data analysis. XL, LL, JY, HC, ZC, SY, LG, LQ and XL conducted the experiments. XW, YL wrote the manuscript. All authors read and approved the final version of the paper.

## DECLARATION OF INTERESTS

The authors declare no competing interests.

**Table S1. Summary of soybean accessions used in this study.**

**Table S2. Phenotypic statistics for symbiotic nitrogen fixation (SNF)-related traits in soybean accessions.**

**Table S3. Gene-to-module annotation based on Independent Component Analysis (ICA).**

**Table S4. Module-level TWAS associations for four SNF-related traits based on ICA components.**

**Table S5. Transcriptome-Wide Association Study (TWAS) results for four SNF-related traits.**

**Table S6. CUT&Tag peaks and genomic annotations in *GmLHY1b*-overexpressing plants.**

**Table S7. Gene expression matrix (CPM) for GmLHY1b overexpression (OE) and control samples (EV).**

**Table S8. Selective sweep signals of lead eSNPs and eGenes.**

**Table S9. Open chromatin regions (OCRs) detected by ATAC-seq in mature nodules (28 DAI) and their overlap with eQTLs.**

**Table S10. eQTLs and regulatory classifications of functionally characterized genes identified among the eGenes.**

---

**Figure S1.**
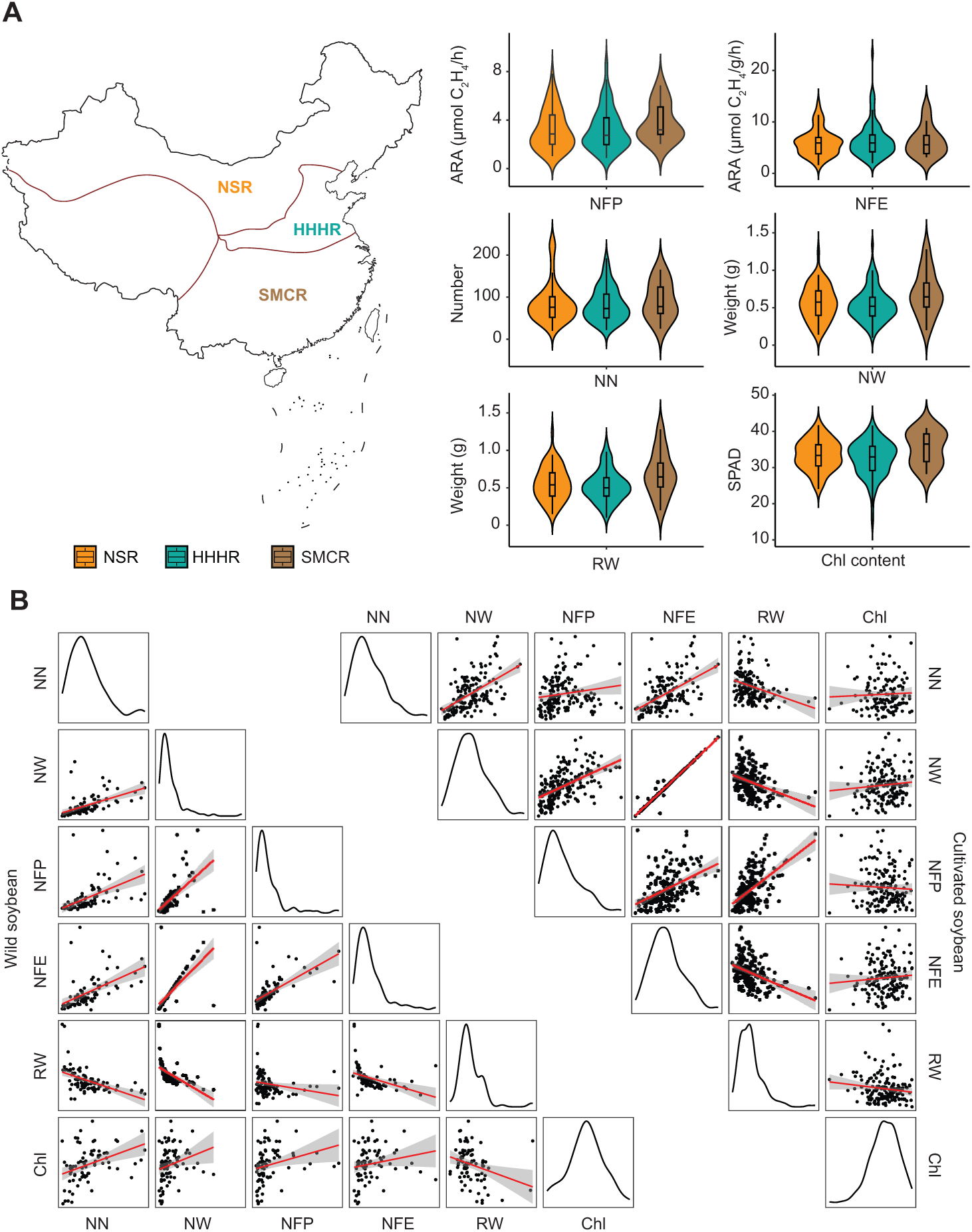
Comparative analysis of six symbiotic traits among accessions from the Huang-Huai-Hai Region (HHHR) and the South–Central Multiple–Cropping Region (SMCR) (a), and the distribution and pairwise correlations of these traits in wild (Glycine soja) and cultivated (Glycine max) soybean accessions (b).

**Figure S2.**
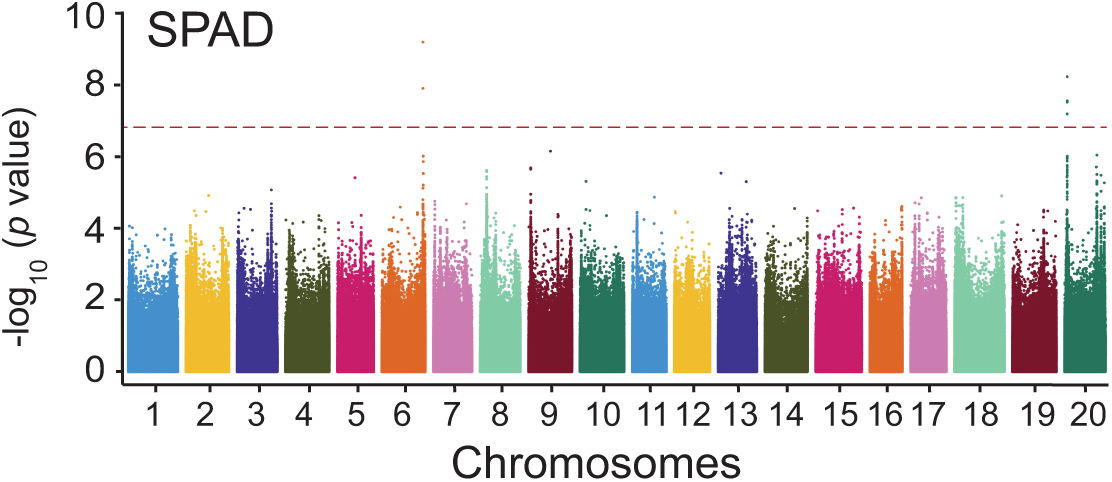
Manhattan plots of GWAS result of SPAD trait. The horizontal dashed line indicates the genome-wide significance threshold (−log10(p) > 6.82).

**Figure S3.**
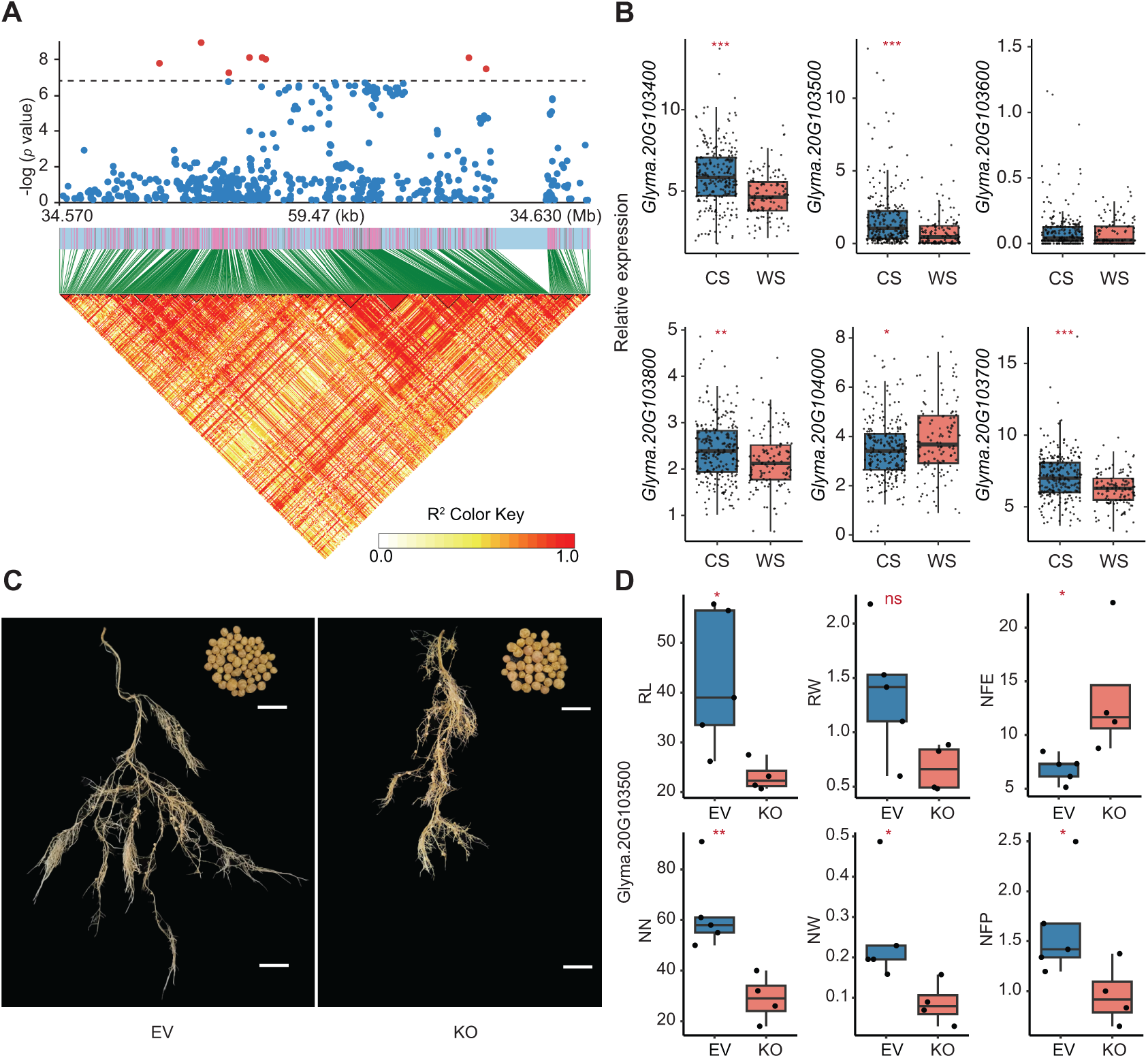
Expression and functional analysis of the candidate region on chromosome 20 associated with nitrogen fixation efficiency (NFE). (A) scatter plot showing SNP association signals within the candidate region on Chr20. Red dots indicate significantly associated loci. (B) Expression patterns of six genes within the candidate region comparing cultivated (CS) and wild soybean (WS) accessions. *P < 0.05, **P < 0.01, ***P < 0.001; ns, not significant. (C) Root and nodule phenotypes at 28 days after inoculation (DAI) following CRISPR-Cas9 hairy root transformation. Scale bar = 1 cm. (D) Quantification of six traits, including root length (RL), nodule number (NN), root weight (RW), nodule weight (NW), nitrogen fixation efficiency (NFE), and nitrogen fixation activity per plant (NFP), in empty vector (EV) and knockout (KO) samples.

**Figure S4.**
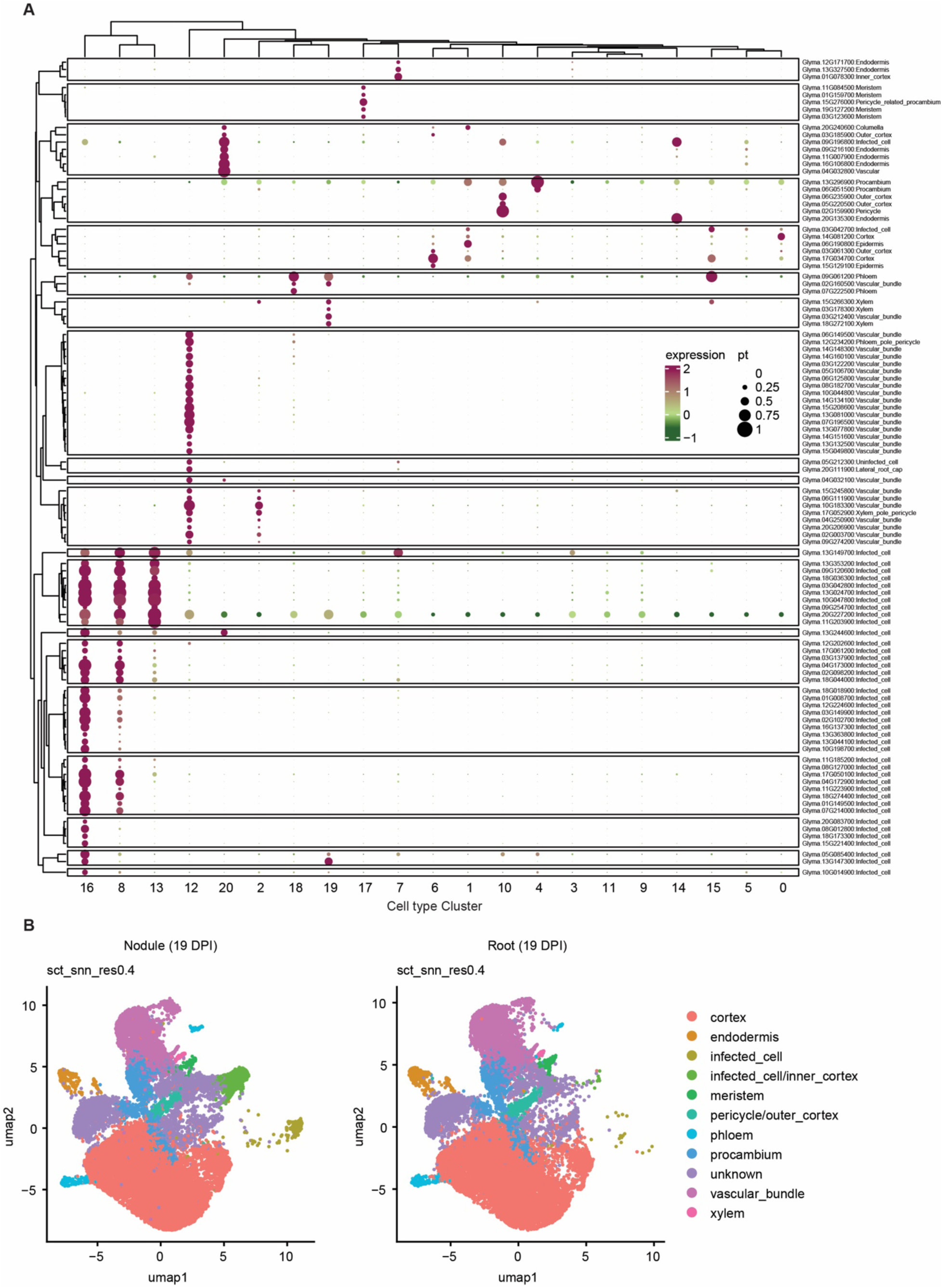
Cell type clustering of mature soybean nodules based on public single-cell RNA sequencing (scRNA-seq) data. (A) Cell type clusters were defined using known marker genes. (B) UMAP visualization of clusters annotated based on marker gene expression. Colors indicate distinct cell type clusters. The scRNA-seq dataset was obtained from public dataset. Briefly, all analyses were performed using the **Seurat** R package. Gene expression matrices from the two species were normalized using the **SCTransform** function. Linear dimensionality reduction was conducted using **RunPCA**, and the top 30 principal components were used for non-linear dimensionality reduction and visualization via **RunUMAP** (parameters: *min.dist* = 0.5, *n.neighbors* = 30). Cell clustering was performed using **FindNeighbors** and **FindClusters**.

